# Influence of food and *Nosema ceranae* infection on the gut microbiota of *Apis cerana* workers

**DOI:** 10.1101/375576

**Authors:** Shao K. Huang, Kun T. Ye, Wei F. Huang, Bi H. Ying, Xin Su, Li H. Lin, Jiang H. Li, Yan P. Chen, Ji L. Li, Xiu L. Bao, Jian Z. Hu

## Abstract

**Background:** Gut microbiota plays an essential role in bee’s health. To elucidate the effect of food and *Nosema ceranae* infection on the gut microbiota of honeybee *Apis cerana,* we used 16S rRNA sequencing to survey the gut microbiota of honeybee workers fed with sugar water or beebread and inoculated with or without *N. ceranae*.

**Results:** The gut microbiota of *A. cerana* is dominated by *Serratia, Snodgrassella,* and *Lactobacillus* genera. The overall gut microbiota diversity was significantly differential by food type. The *N. ceranae* infection significantly affects the gut microbiota only at bees fed with sugar water. Higher abundance of *Lactobacillus, Gluconacetobacter* and *Snodgrassella* and lower abundance of *Serratia* were found in bees fed with beebread than with sugar water. *N. ceranae* infection led to higher abundance of *Snodgrassella* and lower abundance of *Serratia* in sugar-fed bees. Imputed bacterial KEGG pathways showed the significant metagenomics functional differences by feeding and *N. ceranae* infections. Furthermore, *A. cerana* workers fed with sugar water showed lower *N. ceranae* spore loads but higher mortality than those fed with beebread. The cumulative mortality was strongly positive correlated (rho=0.61) with the changes of overall microbiota dissimilarities by *N. ceranae* infection.

**Conclusions:** Both food and *N. ceranae* infection significantly affect the gut microbiota in *A. cerana* workers. Beebread feeding not only provide better nutrition but also help establish a more stabled gut microbiota therefore protect bee in response to *N. ceranae* infection.

**Abstract Importance:** Gut microbiota plays an essential role in bee’s health. Scientific evidence suggests the diet and infection can affect the gut microbiota and modulate the gut health, however the interplay between those two factors and bee gut microbiota is not well known. In this study, we used high-throughput sequencing method to monitor the changes of gut microbiota by both food intake and the *Nosema ceranae* infection. Our result showed that the gut microbiota composition and diversity of Asia Honeybee was significantly associated with both food intake and the *N. ceranae* infection. More interestingly, bees fed with beebread showed higher microbiota stability and less mortality than those fed with sugar water when infected by *N. ceranae*. Those data suggest the potential role of beebread, not only providing better nutrition but also helping establish a more stabled gut microbiota to protect bee against *N. ceranae* infection.

## Background

European honey bees (*Apis mellifera*) and Asian honey bees (*A. cerana*) are two truly domesticated bee species that play a vital role in agriculture and ecosystem by providing pollination service to food crops and natural plants. However, both bee species are confronted with many biotic and abiotic stressors including diseases caused by pathogens and parasites, acute and sublethal toxicity of pesticides, malnutrition due to loss of foraging habitat, and etc that act separately or synergistically to cause the significant decline of bee health and population worldwide[1–3]. As a result, the health of managed honey bees has drawn much attention worldwide in recent years. There has been growing evidence that gut bacteria play very important roles in animal health by maintaining homeostasis, modulating immunity, regulating nutrition metabolism, and supporting host development, and reproduction[4–6]. Although most insect guts harbor relatively few microbiota species as compared to mammalian guts, insect bacteria have been shown to be vital in regulating various aspects of their host biology [7–9]. Over the past decade, progress has been made in understanding the composition and functional capacity of microbes living in honey bee guts [10–12]. Honey bee gut microbiota is established gradually through trophallaxis, food consuming, and interacting with the hive environment[13]. Many factors, like genetics, age, diet, geography, and medication can affect the gut microbiota composition[14,15]. Several types of bacteria have been identified in the guts of *A. mellifera* including the genera of *Bacillus*, *Lactobacilli* and *Staphylococcus* from *Firmicutes* phylum, *Coliforms* from *Enterobacteriaceae* family of *Proteobacteria* phylum [16–18].A previous study reported that species within the *Apis* genus share rather simple and similar gut bacterial microbiota. At phylum level, among *proteobacteria, Gammaproteobacteria* class was the most abundant, while other *proteobacteria* including *Betaproteobacteria* and *Alphaproteobacteria* classes, *Firmicutes* and *Actinobacteria* were less frequent but widespread organisms. Less than ten members formed a core species, including *Lactobacillus, Bifidobacterium, Neisseria, Pasteurella, Gluconobacter* and newly named species: *Snodgrassella* and *Gilliamella* [19–21]. However, most the studies about the microbiota in *Apis* were conducted in European honey bees, *A. mellifera.* The food influence on the microbiota of *A. cerana* has barely been investigated.

*Nosema ceranae* is an intracellular parasite that disrupts a bee’s digestive system. It was first discovered in the *A. cerana* but has recently jumped host from *A. cerana* to *A. mellifera[22,23], N. ceranae* can seriously shorten the life expectancy of adults, decrease the productivity of the colony, and cause severe colony lost especially during wintering in the temperate area[24,25]. Furthermore stresses caused by *Nosema* would be more severe when mixed infection happened with other parasites or pathogens, such as *Varroa* mites, and viruses [26–30]. Now *Nosema* is one of the major threats to the honey bee populations and has been often implemented in honey bee colony losses worldwide[31,32]. The survey of microbial communities from the digestive tracts of *A. cerana* workers showed that *N. ceranae* infection might have detrimental effects on the gut microbiota[1]. However, the relationship between *N. ceranae* and microbiota in *A. cerana* is largely unknown. In this study, we challenged *A. cerana* workers with *N. ceranae,* and then fed them with either beebread or sugar water. The intent of the current study was to evaluate the effects of *N. ceranae* infection and food types on the gut microbiota.

## Methods

### Honey bees

Three *A. cerana* colonies without identified diseases were chosen for sample collection, which located at the campus of College of Bee Science, Fujian Agriculture and Forestry University, Fuzhou, Fujian, China. Capped brood-combs with pupae near emergence were taken out of the colonies and then kept in the incubator with 35±1°C and 55%-65%RH. Workers emerged within 24h were collected for the study.

#### Purification of *Nosema ceranae* spore

Because the prevalence and spore loads of *N. ceranae* in *A. cerana* are less than *A. mellifera[33,34],* we purified *N. ceranae* spores from *A. mellifera* foragers. First, adult workers were captured at entrances of *A. mellifera* colonies and immobilized in the refrigerator for few minutes, and then the guts of the bees were dissected, pooled, and ground in a mortar. Afterward, the spores were purified by differential centrifugation to exclude most of the debris, finally, the suspension was loaded on Percoll (Sigma-Aldrich, St.Louis, USA) and centrifuged to eliminate unsaturated spores [33]. The purity and maturity of spore were confirmed under phase contrast microscopy. The *Nosema* species was confirmed by PCR method [35].

### Treatments and sampling

The newly emerged workers (<24h) were randomly distributed into 18 laboratory rearing cages. 30 bees were transferred to each cage The experimental cages were divided into two groups: 1) group supplied with only 50% (W/V) sugar water in modified syringe feeder [36], and 2) group supplied with both 50% (W/V) sugar water and beebread freshly collected from the *A. cerana* colonies (thereafter call beebread). For each group, three subgroups were set up one without spore inoculation which was used as a negative control, one inoculated with *N. ceranae* 5000 spores per bee, and one inoculated with 50000 spores per bee (Figure 1). Each subgroup consisted of three cages as replicates. Cages were kept in an incubator with 30±1°C and 55%-65% RH. About eight workers were collected at day 5, 10, and 15 post treatment (dpi) from each subgroup. The gut tissue was collected from each bee at 5-day, 10-day, and 15-day post infection and then stored into −80°C freezer until the subsequent microbial composition analysis Foods were changed each other day; dead bees were counted and removed every day.

**Figure 1.**
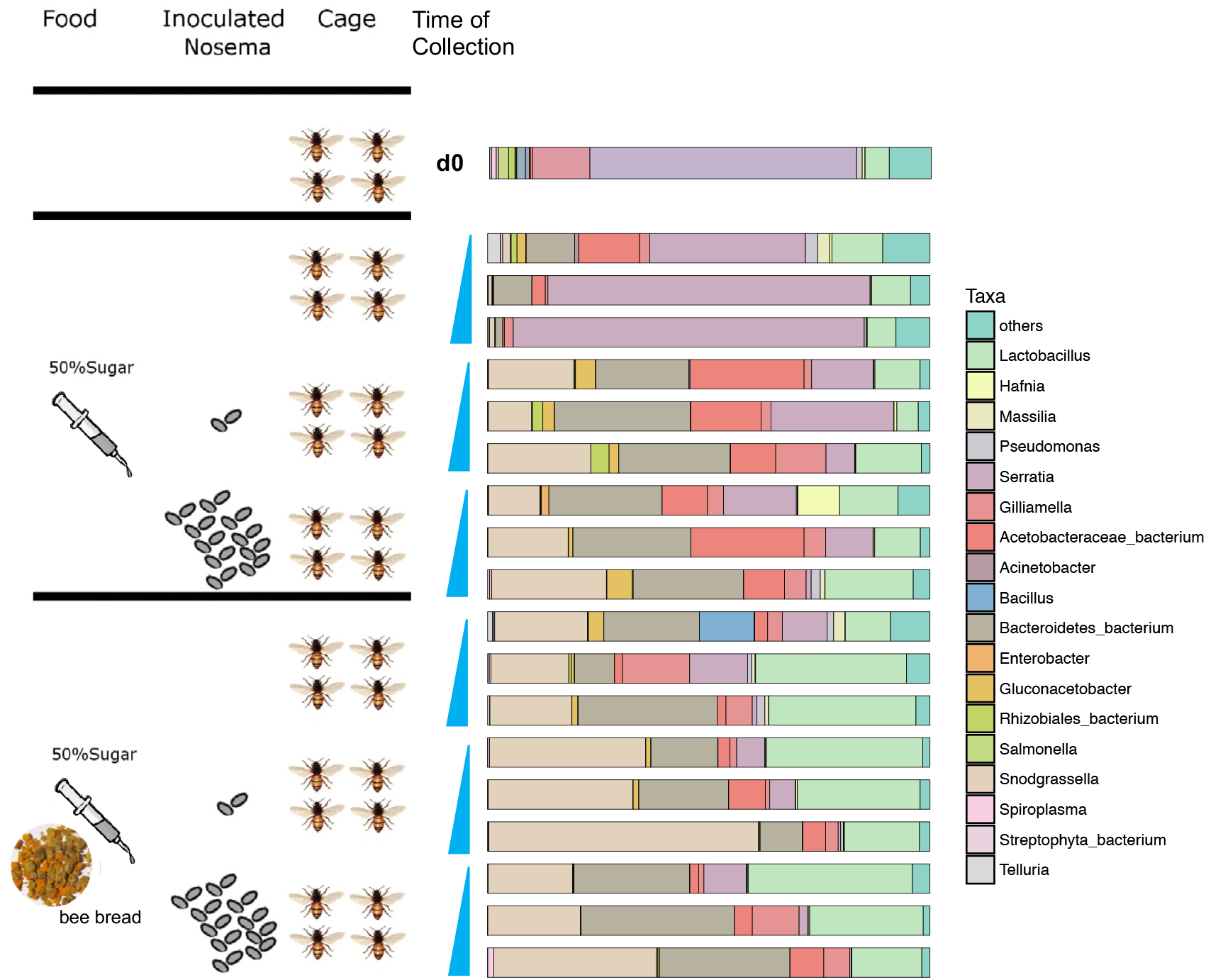
Experimental design and survey of the gut microbiota composition. Bar plots represent the normalized relative abundance (%) of the gut microbiota at genus level fed with different foods and doses of *N. ceranae* infection at day 5, 10 and 15 (increasing order).

### DNA extraction from gut tissue samples

Sample bees were taken out of the refrigerator and rinsed with 7% benzalkonium bromide for 2min and then rinsed four times with sterilized water to minimize the bacterial contamination from the body surface. The intestine tissues were collected with tweezers clamping the end of the abdomen and each gut tissue was further separated and transferred into a labeled 1.5ml tube on ice. The entire procedure was conducted under the aseptic condition and all tools used were sterilized. The total DNA of the gut tissue samples was extracted using Insect DNA Extraction Kit II (Beijing Demeter Biotech Ltd, Beijing, China) following the manufacturer’s instruction. The quality and yield of DNA samples were assessed using a Quawell Q5000 UV-Vis spectrophotometer (Quawell, San Jose, CA, USA).

### Gut *Nosema ceranae* spore counting

After caged bees were sampled at day 5, 10, and 15 post treatment, the quantity of the spores in the gut specimen was counted as previously described with slight modification[33]. Briefly, the sediments of gut were resuspended in 100μl ddH2O, then vortexed evenly. The suspension was loaded onto the hemocytometer for *N. ceranae* spore inspection and counting under a microscope. We conducted three to four repeated measurements for each sample.

### Bacterial 16S ribosomal RNA gene PCR amplification

The phylogenetically informative V3-V4 region of 16S ribosomal RNA (rRNA) gene was amplified using universal primer 347F/803R [37]. The dual-barcoding approach as previously described[38] was applied to label the 16S rRNA gene amplicons of each sample. Briefly, the 6-mer barcodes were attached on the 5’ends of both forward and reverse PCR primers so that 16S rRNA gene PCR amplicons from each sample contained a unique dual barcode combination. The PCR Primers were synthesized by Sangon Biotech, Shanghai, China, and the primer sequences are shown in Supplementary Table 1. The 25-μL PCR reaction mixes contain 300ng of sample DNA as PCR template, 1μL of 10μM forward and reverse 16S primers, and 12.5μL of 2×HotMaster Taq DNA mix (Tiangen Biotech, Beijing, China). The PCR reaction was performed on Applied Biosystem 2720 thermal cycler (Thermo Fisher Scientific Inc., Waltham, MA, USA) at 94°C for 3 minutes, then 94°C 30 seconds, 58°C 30 seconds, and 72°C 20sec for 30cycles, and 72°C for 4 min. The integrity of the PCR products was verified by agarose gel electrophoresis. After purified with gel purification kit (Promega, Madison, WI, USA), the 16S PCR amplicons were pooled at equal molarity, freeze-dried, and submitted to New York Genome Center for sequencing.

### 16S rRNA gene sequencing and microbiota profiling

The 16S rRNA gene PCR amplicons were sequenced on the Illumina HiSeq platform using 2×250 paired-end fast-run mode. In total, we generated 21 million high-quality 16S reads obtained by NGS sequencing on pooled barcoded PCR amplicons from 86 samples. After splitting by barcodes, ~ 2.5×10^5^ reads per sample were obtained. After the merge, the sequencing reads with length >400 and the quality score >Q30 at more than 99% of bases were further split by barcode and trimmed of primer regions using CLC Genomic workbench 6 (Qiagen Bioinformatics, Redwood City, CA, USA). The filtered and trimmed high-quality reads were further processed by QIIME 1.9.0[39]. We used the command *pick_open_reference_otus.py* with the defaulted cutoff =97% to a cluster of nearly-identical sequencing reads as an Operational Taxonomic Unit (OTU) using *Uclust*[40]. Representative sequences for each OTU were aligned using PyNAST. Finally, the program built a biom-formatted OTU table with assigned taxonomical information for each OTUs. Using Chimera Slayer[41], chimera sequences arising from the PCR amplification were detected and excluded from the aligned representative sequences and the OTU table.

### Statistical Analysis

The mortality data of different groups were transformed by square root and degrees and Asin, and then compared by using two-way ANOVA of the SPSS program. The overall microbiota dissimilarities among all samples were accessed using the Bray-Curtis distance matrices[42] generated at the genus level. The PERMANOVA (Permutational Multivariate Analysis of Variance) procedure [43,44] using the [Adonis] function of the *R* package *vegan* 2.0-5), with the maximum number of permutations = 999, was performed to test the significance of the overall microbiota differences between the gut microbiota grouped by feeding types and *N. ceranae* infections. The diversity within each microbial community, so-called alpha-diversity was calculated using the Shannon Index as metric and represented the measure of the diversity at genus level [45]. Using the linear discriminant analysis (LDA) effect size (LEfSe) method[46], we further selected the microbiota features significantly associated with feeding types and *N. ceranae* infections. The program PICRUSt (Phylogenetic Investigation of Communities by Reconstruction of Unobserved States)[47] was used to predict the metagenome functional content based on our 16S rRNA gene sequencing data. Briefly, a close reference-based OTU table was generated using the QIIME pipeline and input into PICRUSt to bin individual bacterial genes into Kyoto Encyclopedia of Genes and Genomes (KEGG) pathways, thereby predicting their function.

### Dataset

16S rRNA gene sequencing information has been deposited in the European Nucleotide Archive with study accession number: PRJEB21090.

## Results

### 1. Simple core bacterial clusters in the gut of *Apis cerana*

As illustrated in Figure 1, the *A. cerana* adult workers were grouped by foods and the level of *N. ceranae* infection. The microbial composition analysis of gut tissue collected at 5-day, 10-day and 15-day post infection for each subgroup following the method described previously [18,49] showed that the gut microbiota of *A. cerana* is rather simple and mainly contain three phyla, *Proteobacteria*, *Firmicutes*, and *Bacteroidetes*, counting for over 97% of the total microbiota composition (Figure 1, Figure 2). At the genus level, less than 6 taxa from *Proteobacteria* and *Firmicutes* are dominant in the *A. cerana* gut bacterial community. In details, they were the genera *Snodgrassella, Acetobacteraceae, Serratia, Gilliamella, Lactobacillus* and unclassified genus from *Bacteroidetes*, of which *Serratia* was not in the core species clusters of *A. mellifera*[50].

**Figure 2.**
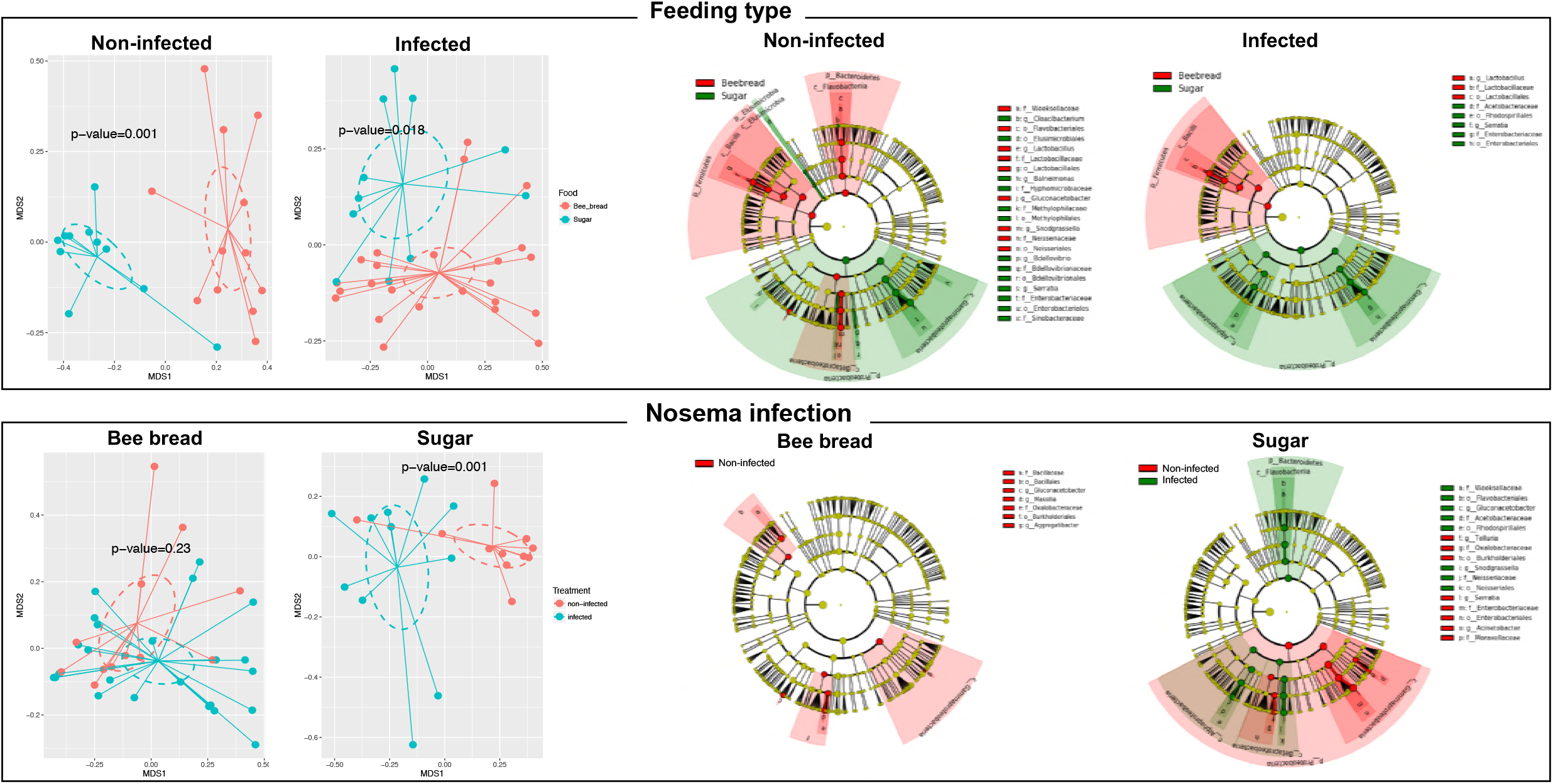
Foods and *N. ceranae* infection changed the relative abundance of microbes in the gut. 2A. NMDS plots present the overall dissimilarity between samples grouped by diet or infection status. *P*-values were given by PERMANOVA test. 2B. The cladogram plots present the LEfSe results on the gut microbiota of honey bees grouped by diet or infection status. Differences are represented in the color for the most abundance class (green and red color indicate increasing gin corresponded phenotype).

### 2. Foods and *N. ceranae* infection changed the relative abundance of microbes in the gut

The overall microbiota dissimilarity in samples grouped by food or *Nosema* infection was visualized in NMDS plots (Figure2). The overall gut microbiota is significant different between bees fed with beebread fed and sugar (p=0.018 with *N. ceranae* infection, and p=0.001 without infection, PERMANOVA test using Bray-Curtis distance). In sugar fed bees, we found *N. ceranae* infection significantly altered the microbiota (p=0.001). However, *N. ceranae* infection caused no significant alteration in gut microbitota in bees fed with beebread (p=0.23). LEfSe method was applied to select the microbiota taxa which are significantly associated with either food types or *N. ceranae* infections. In subgroups without *N. ceranae* infection, the bees fed with beebread showed more abundant *Lactobacillus, Snodgrassella, Weeksellaceae,* and less abundant *Serratia* genus than bees fed with sugar. However, in the subgroups with *N. ceranae* infection, the bees fed with beebread showed more abundant OTUs of *Lactobacillus* and less abundant *Serratia* and *Acetobacteraceae* than bees fed with sugar (Figure2). Among bees fed with sugar solution, *N. ceranae* infection caused major changes in microbiota and was associated to increased OTUs of *Weeksellaceae, Snodgrassella and Gluconacetobacter* and decreased *Proteobacteria* phyla, in particular, *Telluria, Serratia,* and *Acinetobacter*. Among bees fed with beebread, *N. ceranae* infection had a minor effect on microbiota, with merely decreased the abundance of *Massilia, Aggregatibacter* and *Gluconacetobacter* genera.

### 3. Differential metagenome features predicted by PICRUSt and their association with food and *N. ceranae* infection status

We performed PICRUSt analysis to predict the full metagenomic content of microbial communities using 16S gene surveys^33^ and compared the predicted metagenomic pathways by food and *N. ceranae* infection status (FigureS1). The Nearest Sequenced Taxon Index (NSTI), which quantifies the uncertainty of the prediction (lower values mean a better prediction), ranged from 0.027 to 0.11 with mean value=0.067, indicating fair reliability and accuracy in the metagenome reconstruction. The heat map (FigureS1) with clustering analysis showed the overall changes in predicted KEGG pathways. Among those significantly differential pathways, we found the food type could affect the bacterial Glycolysis/Gluconeogenesis, Fructose and mannose metabolism, metabolism of several amino acids and etc. *N. ceranae* infection could affect biosynthesis of several amino acids, the signal transduction mechanism, and the lipopolysaccharide biosynthesis and phosphotransferase system (PTS).

### 4. The cumulative mortality of caged bees with different feeding type and infection status

When inoculated with *N. ceranae* spores, the average cumulative mortality of caged bees increased gradually during the experimental observation, our results showed that *N. ceranae* infection significantly shortened the longevity of workers fed with only sugar water than those fed with beebread (Figure 3A). Interestingly, the spore load in the gut fed with beebread were significantly higher than those with sugar water (p-value=0.01 and 0.007 for low *N. ceranae* and high *N. ceranae,* respectively) at 15 days after inoculation (Figure 3B). This was consistent with the earlier report by Zheng et al[51]. There is not interaction between food type and spore dosage on the mortality (p-value=0.868, F=0.029). There was no significant difference in gut *N. ceranae* spore counts between low and high dosage *N. ceranae* inoculations. This may be due to the late sampling time that the spore load in the gut has reached the plateau.

**Figure 3.**
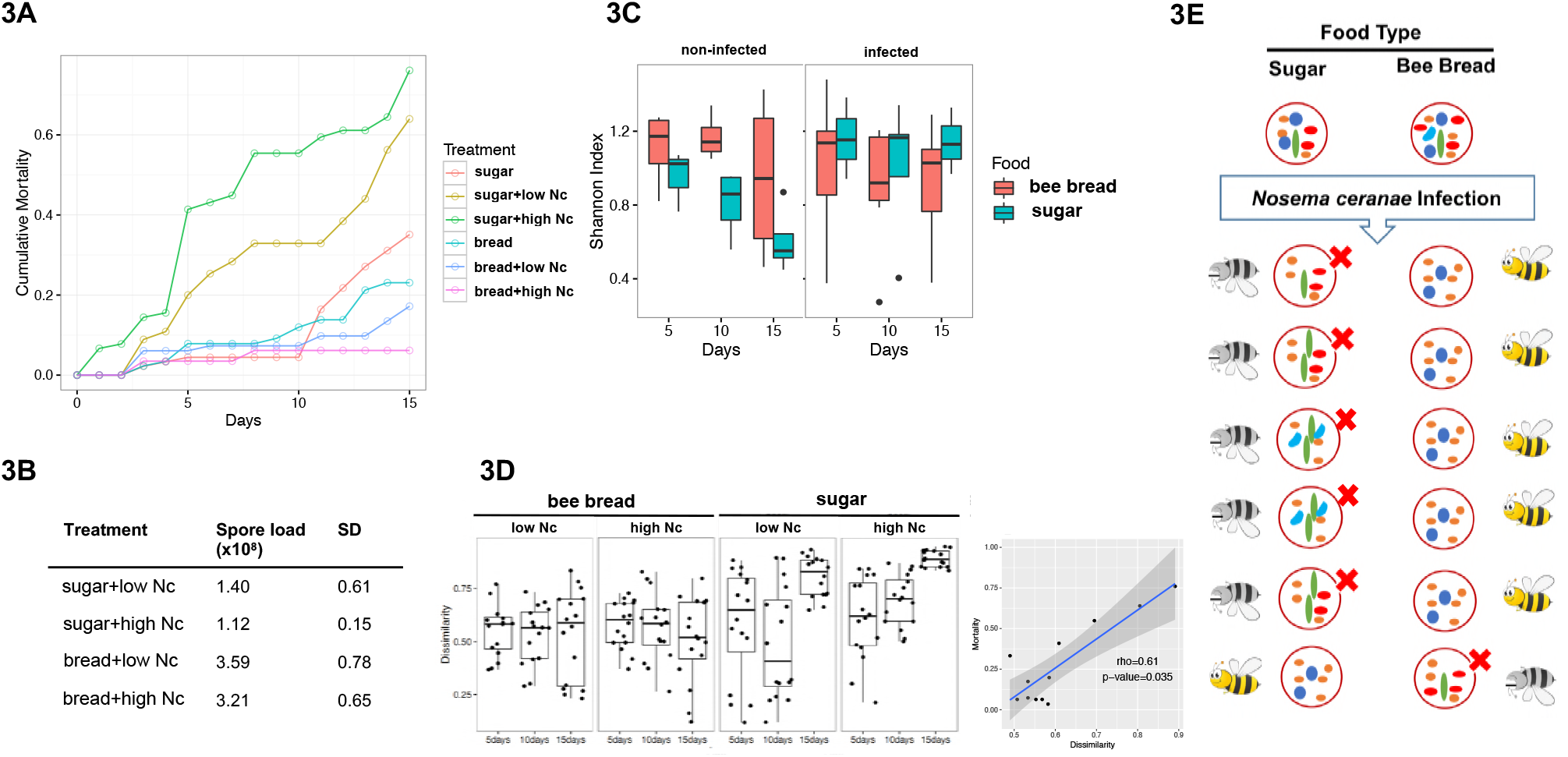
Difference in bee mortality and stabilities of the gut microbiota by feeding and *N. ceranae* infections. 3A.Cumulative mortalities under different treatment conditions; 3B. The mean and standard deviations of the *N. ceranae*(Nc) spore load in different treatment conditions; 3C. Alpha diversity of the mid-gut microbiota in different treatment conditions; 3D. The mean and variance of the dissimilates (Beta diversity) of the mid-gut microbiota in different treatment conditions; 3E. Illustration of the links between the decreased stability in mid-gut microbiota and the increased bee mortality in sugar-fed bees.

### 5. The richness of the gut microbiota of caged bees with different feeding type and infection status

We used Shannon index, a commonly used metrics, for richness assessment within the given community[45]. Without *N. ceranae* infection, the richness of the microbiota in bees fed with sugar solution was significantly lower than those fed with beebread (Figure 3C, p-values<0.05 at 5, 10 and 15days). Furthermore, the richness of the microbiota decreased by time in bees fed with sugar, but not in bees fed with beebread. In subgroups with *N. ceranae* infection, the microbiota of bees fed with sugar showed increased richness at all time points with a slightly higher mean but no significant differences to that of bee fed with beebread.

### 6. The stability of the gut microbiota is significantly correlated with the cumulative mortality

The stability of the microbiota in response to the *N. ceranae* infection in groups with different of feeding conditions showed that the bee fed with beebread showed relatively stable microbiota. The mean dissimilarity was not significantly different from either sampling time post infection or *N. ceranae* doses. However, among bees fed with sugar solution, the microbiota dissimilarities significantly increased by time, with the most dissimilarities and higher consistency at 15 days with the high *N. ceranae* infection (Figure 3D), suggesting the most diverged microbiota within this group. Further, the mean microbiota dissimilarities were significantly correlated with the cumulative mortality rate (r=0.61, spearman correlations, p-value=0.035).

## Discussion

Our result demonstrated that the gut microbiota of the *A. cerana* adult workers are composed of three major phyla, *Proteobacteria, Firmicutes,* and *Bacteriodetes.* This result is consistent with the previous reports [21] except that the most abundant taxa in our study was *Proteobacteria,* which was the second in Ahn’s study [21]. At the genus level, we found that the gut microbiota of Asian honey bees is dominated by a few core bacterial species, including *Lactobacillus*, *Snodgrassella* and *Gilliamella,* among the major genera found in both our study and previous studies[52].

Food constituents can influence the gut microbiota composition. Our results confirmed that food type significantly shapes the bees’ gut microbiota composition (Figure 1 and 2). Beebread contains high protein and comprehensive nutrients, which may favor those proteolytic species. In addition, beebread provides additional microbiota [53,54] inoculations especially lactic acid bacteria, and may benefit the gut microflora too.

Our study also showed that *Lactobacillus* and *Snodgrassella* genera were much more abundant in those bees fed on beebread (Fig 2B). The genera, *Lactobacillus, Bifidobacterium* and the family *Pasteurelaceae,* were also found in beebread from colonies of *A. mellifera* [51]. *Lactobacillus* had been found in flora and hive environment, including honey, royal jelly, beebread, and honey sac. *Lactobacillus* was also found in honey bee crop and showed inhibition effect on *Paenibacillus larvae* in vitro [55]. Therefore, it is plausible to speculate that the *Lactobacillus* found in gut of adult workers fed with were obtained through food trophallaxis. In contrast, bees fed with sugar only showed more abundant *Enterobacteraceae*. Overgrowth of *Enterobacteraceae* has been linked to gut inflammation in many studies.

Our data showed that higher proportion of *Serratia* harbored in the gut of 10-day old bees fed with sugar. *Serratia* was further confirmed as *S. marcescens* by sequencing near full-length 16S rRNA gene(data no showed). *S. marcescens* is commonly found in adult *A. mellifera, A. cerana,* and bumble bee gut. It is generally harmless to honey bee, and commonly used to explore the host immune reaction to microbes[56], There were two cases, that *Serratia* had detrimental effects on *A. mellifera* survivorship after host microbiota was erased by antibiotics. In our study, *N. ceranae* infection broke the balance of *Serratia* in the microflora, and shortened host lifespan. Future investigations are necessary to further explore complex interactions among *N. ceranae,* host, and gut microbiotas.

Nosema resides in the gut of the bee and the infection by *N. ceranae* can profoundly change honey bees physiology [57], and change the host-microbiota relationship in the gut. Investigation conducted by Li et al. showed that four common bacterial clusters, *Bifidobacterium, Neisseriaceae, Pasteurellaceae,* and *Lactobacillus* in *N. ceranae* infected adult *A. cerana* workers were less abundant compared to non-infected ones[58]. However, we found minor changes in gut microbiota by *N. ceranae* infection in beebread fed bees. When sugar water is the only food supplied, *N. ceranae* infection showed a stronger effect on the overall gut microbiota with more abundant *Neisseriaceae/Snodgrassella, Weeksellaceae, Gluconacetobacter* and less abundant *Serratia, Telluria* and *Enterobacteriaceae*. Further, the lower stability of gut microbiota in bee fed with sugar could lead to increased susceptibility to Nosema infections in bees.

Our data showed that the *N. ceranae* infection caused much higher cumulative mortality in bees fed with sugar than bees fed with beebread. Interestingly, the changes in microbiota dissimilarity were highly correlated to bee’s mortality. *N. ceranae* infection caused significant increases in both the microbiota richness and the dissimilarity in sugar fed bees, but not beebread fed bees. We speculated that *N. ceranae* infection in sugar fed bees resulted in a more diverged microbiota, among which many are not considered as probiotic in bees. The gut microbiota in bees fed with beebread was stable with *N. ceranae* infection. This stability of gut microbiota could play a protective role and result in less mortality.

Having a biological measure of the effect of *N. ceranae* infection might help us further understand the controversy of honey bee health and *N. ceranae* infection, which bees with pollen feeding resulted in higher spore load but less mortality compared to those with sugar water[59].

Although the 16S sequencing based taxonomy analysis is sufficient in current technology development, it only identified bacterial taxa to genus level. It is difficult to identify a specific species or strain that is strongly correlated to either the food feeding or *N. ceranae* infection.

In summary, the gut microbiota of *A. cerana* workers is significantly differentiated by both food types and *N. ceranae* infection. The higher stability of the gut microbiota in the bees fed with plays a role in bees ability to defend *N. ceranae* infection and warrants further exploration

## Authors’ contributions

SKH and JZH conceived an designed the study; KTY and SKH performed the bee experiment, KTY, BHY, XS, and LHL counted spore loads; JZH and XLB performed the microbiota analysis; WFH, JHL and YPC analyzed the phenotype data; SKH, JZH and JLL contributed reagents/materials/analysis tools; SKH, JZH wrote the paper; WFH and YPC revised the paper. All authors read and approved the final manuscript.

## Acknowledgement

We thank the genomic center of NYU to perform the NGS. This project was sponsored by the Scientific Research Foundation for the Returned Overseas Chinese Scholars, State Education Ministry (K4115005A); The Ministry of Agriculture 948 project (2015-Z9).

---

**Supplementary Table 1.**
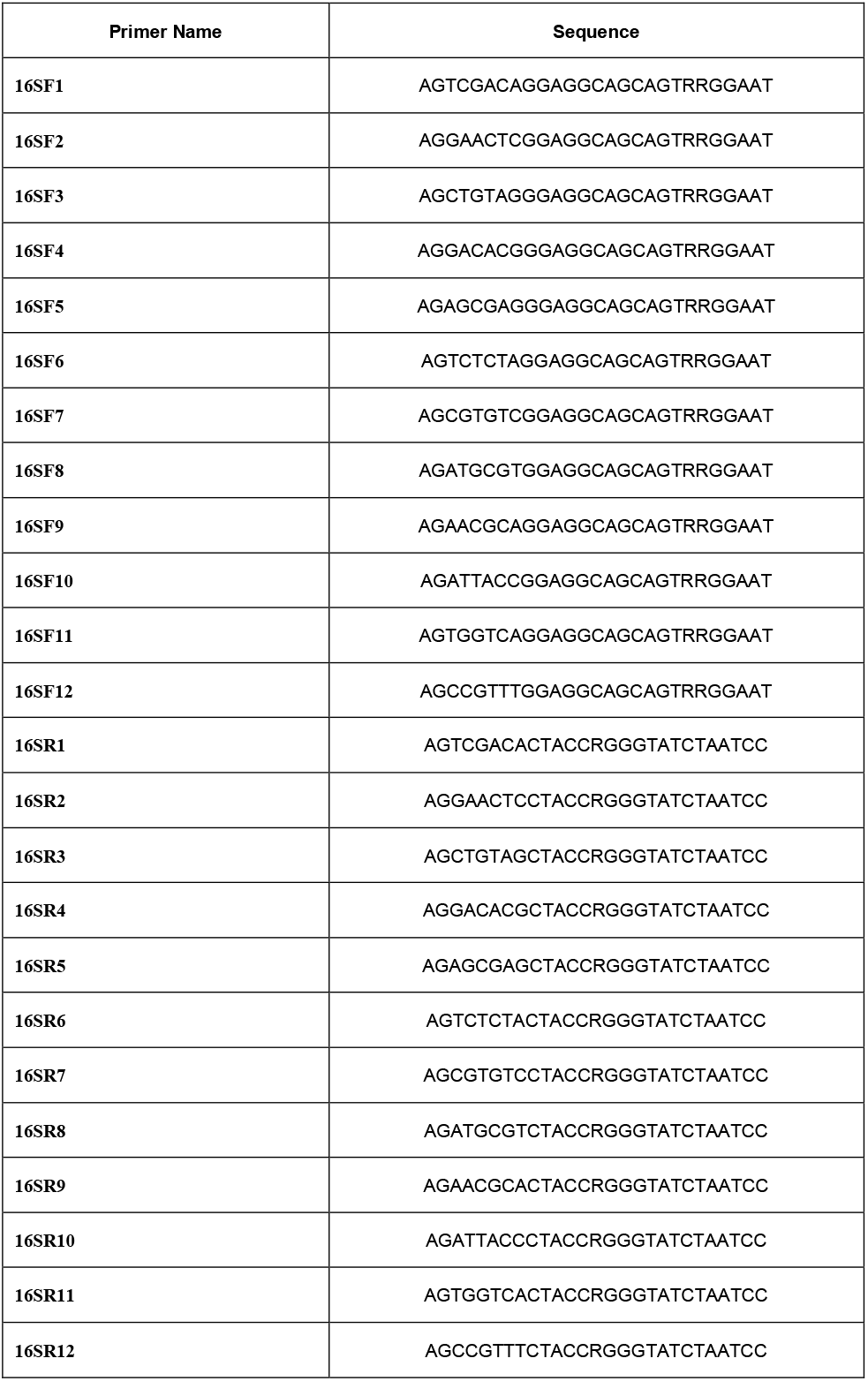
PCR primers for bacterial 16S sequencing

## Supplementary Figures

**Figure S1.**
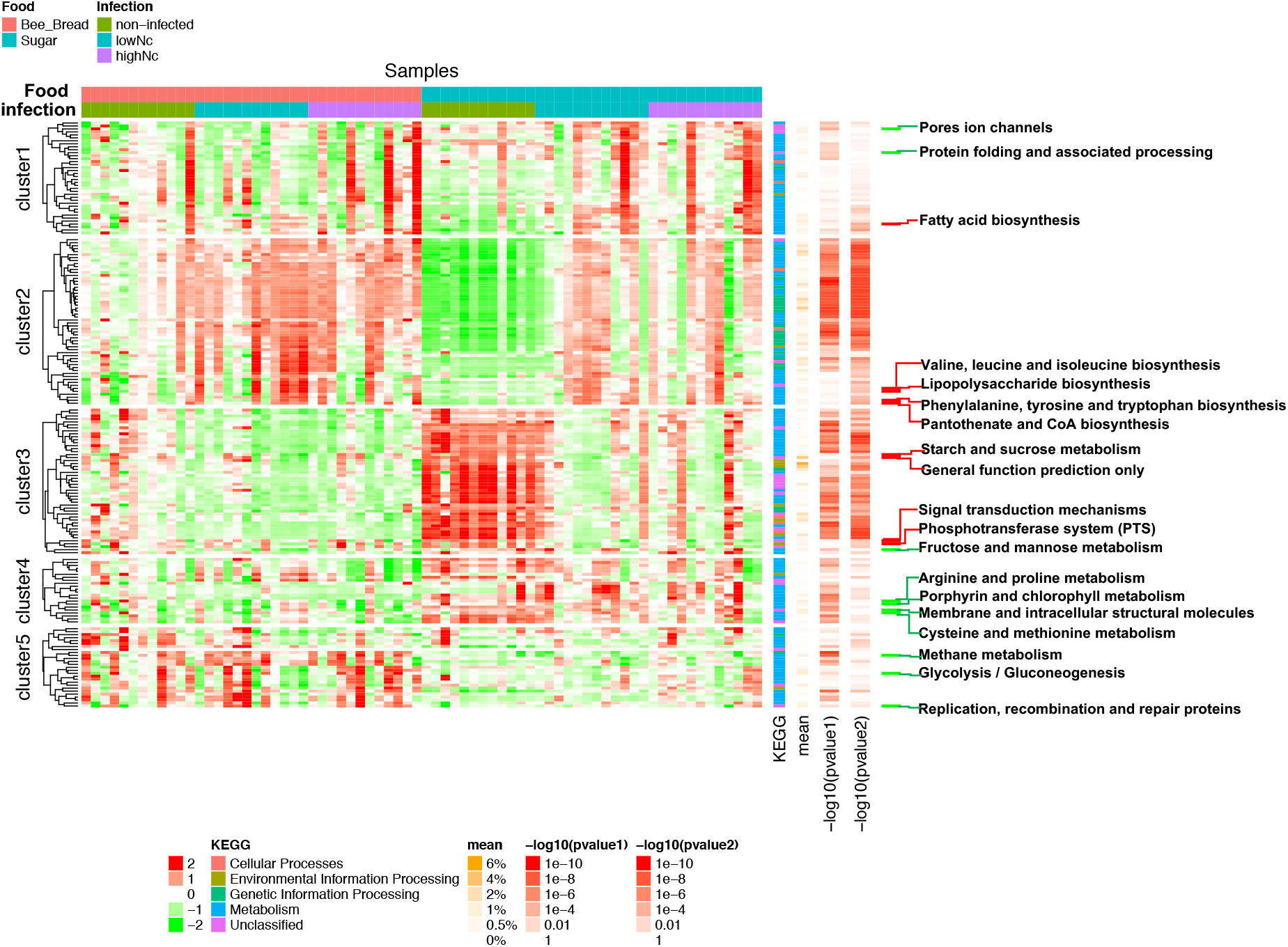
Heatmap of metagenome features predicted by PICRUSt and their association with food and *N. ceranae* infection. Heatmap was drawn by R package *ComplexHeatmap*. Each column corresponds to a specific sample, each row to a KEGG pathway predicted by PICRUSt. The proportions that each lineage contributed to the full population within each sample are indicated with the color scale to the right of the figure (values from −2 to 2). Metadata is color-coded at the top, including food, and *N. ceranae* infection. The KEGG pathways were clustered using average linkage hierarchical clustering as default and split by kmeans=5. The mean abundance of each pathway ranged from 0% to 6% was shown with gradual color changes. A non-parametric Wilcox test with FDA adjusted p-values was listed for food(pvalue1) or infection(pvalue2). The labeled KEGG Pathways are with >0.5% mean abundance and only shown significance(p<0.05) at either food (green color) or infection (red color).

## References

1. Li J, Qin H, Wu J, Sadd BM, Wang X, et al. (2012) The prevalence of parasites and pathogens in Asian honeybees Apis cerana in China. PLoS One 7: e47955.

2. Becher MA, Osborne JL, Thorbek P, Kennedy PJ, Grimm V (2013) Towards a systems approach for understanding honeybee decline: a stocktaking and synthesis of existing models. J Appl Ecol 50: 868–880.

3. Godfray HC, Blacquiere T, Field LM, Hails RS, Petrokofsky G, et al. (2014) A restatement of the natural science evidence base concerning neonicotinoid insecticides and insect pollinators. Proc Biol Sci 281.

4. Wu GD, Chen J, Hoffmann C, Bittinger K, Chen YY, et al. (2011) Linking Long-Term Dietary Patterns with Gut Microbial Enterotypes. Science 334: 105–108.

5. Cantas L, Sørby JR, Aleström P, Sørum H (2012) Culturable gut microbiota diversity in zebrafish. Zebrafish 9: 26–37.

6. Lee WJ, Hase K (2014) Gut microbiota-generated metabolites in animal health and disease. Nature Chemical Biology 10: 416–424.

7. Anderson KE, Sheehan TH, Eckholm BJ, Mott BM, Degrandi-Hoffman G (2011) An emerging paradigm of colony health: microbial balance of the honey bee and hive (Apis mellifera). Insectes Sociaux 58: 431–444.

8. Engel P, Martinson VG, Moran NA (2012) Functional diversity within the simple gut microbiota of the honey bee. Proceedings of the National Academy of Sciences 109: 11002–11007.

9. Pernice M, Simpson SJ, Ponton F (2014) Towards an integrated understanding of gut microbiota using insects as model systems. Journal of Insect Physiology 69: 12–18.

10. Babendreier D, Joller D, Romeis J, Bigler F, Widmer F (2007) Bacterial community structures in honeybee intestines and their response to two insecticidal proteins. FEMS Microbiol Ecol 59: 600–610.

11. Cox-Foster D, Conlan S, Holmes E, Palacios G, Evans J, et al. (2007) A Metagenomic Survey of Microbes in Honey Bee Colony Collapse Disorder. Science 318: 283–287.

12. Kapheim KM, Rao VD, Yeoman CJ, Wilson BA, White BA, et al. (2015) Caste-Specific Differences in Hindgut Microbial Communities of Honey Bees (Apis mellifera). Plos One 10: e0123911.

13. Je P, VG M, K U, NA M (2014) Routes of Acquisition of the Gut Microbiota of the Honey Bee Apis mellifera. Applied and Environmental Microbiology 80: 7378–7387.

14. Kwong W, Engel P, Koch H, Moran N. Genome sequencing reveals host specialization in bee gut symbionts; 2014. IUSSI.

15. Moran NA (2015) Genomics of the honey bee microbiome. Current Opinion in Insect Science 148: 22–28.

16. Kacaniova M, Chlebo R, Kopernicky M, Trakovicka A (2004) Microflora of the honeybee gastrointestinal tract. Folia Microbiol (Praha) 49: 169–171.

17. Jeyaprakash A, Hoy MA, Allsopp MH (2003) Bacterial diversity in worker adults of Apis mellifera capensis and Apis mellifera scutellata (Insecta: Hymenoptera) assessed using 16S rRNA sequences. J Invertebr Pathol 84: 96–103.

18. Mohr KI, Tebbe CC (2006) Diversity and phylotype consistency of bacteria in the guts of three bee species (Apoidea) at an oilseed rape field. Environmental Microbiology 8: 258–272.

19. Kwong WK (2015) Evolution and Specialization of the Gut Microbiota of Eusocial Bees. Dissertations & Theses - Gradworks.

20. Kwong WK, Moran NA (2015) Evolution of host specialization in gut microbes: the bee gut as a model. Gut Microbes 6: 214.

21. Ahn JH, Hong IP, Bok JI, Kim BY, Song J, et al. (2012) Pyrosequencing analysis of the bacterial communities in the guts of honey bees Apis cerana and Apis mellifera in Korea. Journal of Microbiology 50: 735–745.

22. Huang WF, Jiang JH, Chen YW, Wang CH (2007) A Nosema ceranae isolate from the honeybee Apis mellifera. Apidologie 38: 30–37.

23. Fries I (2010) Nosema ceranae in European honey bees (Apis mellifera). J Invertebr Pathol 103 Suppl 1: S73–79.

24. Higes M, Martin R, Meana A (2006) Nosema ceranae, a new microsporidian parasite in honeybees in Europe. J Invertebr Pathol 92: 93–95.

25. Chen YP, Huang ZY (2010) Nosema ceranae, a newly identified pathogen of Apis mellifera in the USA and Asia. Apidologie 41: 364–374.

26. Goblirsch M, Huang ZY, Spivak M (2013) Physiological and behavioral changes in honey bees (Apis mellifera) induced by Nosema ceranae infection. PLoS One 8: e58165.

27. Anderson DL, Giacon H (1992) Reduced Pollen Collection by Honey Bee (Hymenoptera: Apidae) Colonies Infected with Nosema apis and Sacbrood Virus. Journal of Economic Entomology 85: 47–51(45).

28. Toplak I, Jamnikar Ciglenecki U, Aronstein K, Gregorc A (2013) Chronic bee paralysis virus and Nosema ceranae experimental co-infection of winter honey bee workers (Apis mellifera L.). Viruses 5: 2282–2297.

29. Little CM, Shutler D, Williams GR (2016) Associations among Nosema spp. fungi, Varroa destructor mites, and chemical treatments in honey bees, Apis mellifera. Journal of Apicultural Research: 1–8.

30. Bromenshenk JJ, Henderson CB, Wick CH, Stanford MF, Zulich AW, et al. (2010) Iridovirus and microsporidian linked to honey bee colony decline. PLoS One 5: e13181.

31. Jack CJ, Lucas HM, Webster TC, Sagili RR (2016) Colony Level Prevalence and Intensity of Nosema ceranae in Honey Bees (Apis mellifera L.). Plos One 11: e0163522.

32. Emsen B, Guzman-Novoa E, Hamiduzzaman MM, Eccles L, Lacey B, et al. (2016) Higher prevalence and levels of Nosema ceranae than Nosema apis infections in Canadian honey bee colonies. Parasitol Res 115: 175–181.

33. Huang SK, Yang SS, Wang LH, Fu ZM (2007) Infectivity of microsporidium from Apis cerana cerana to Italian honey bee worker. Apiculture of China 58: 7-8,12.

34. Huang SK, Dong J, Zhang ZR, Hong ST, Chao M (2006) Natural infection rate of Nosema in forager workers of Chinese honey bee and Italian honey bee. Journal of Bee.

35. Chen Y, Evans JD, Zhou L, Boncristiani H, Kimura K, et al. (2009) Asymmetrical coexistence of Nosema ceranae and Nosema apis in honey bees. J Invertebr Pathol 101: 204–209.

36. Huang SK, Csaki T, Doublet V, Dussaubat C, Evans JD, et al. (2015) Evaluation of Cage Designs and Feeding Regimes for Honey Bee (Hymenoptera: Apidae) Laboratory Experiments. Journal of Economic Entomology 107: 54–62.

37. Ahn J, Yang L, Paster BJ, Ganly I, Morris L, et al. (2011) Oral microbiome profiles: 16S rRNA pyrosequencing and microarray assay comparison. Plos One 6: e22788–e22788.

38. Torres J, Bao X, Goel A, Colombel JF, Pekow J, et al. (2016) The features of mucosa-associated microbiota in primary sclerosing cholangitis. Alimentary Pharmacology & Therapeutics 43.

39. Caporaso JG, Kuczynski J, Stombaugh J, Bittinger K, Bushman FD, et al. (2010) QIIME allows analysis of high-throughput community sequencing data. Nature Methods 7: 335–336.

40. Edgar RC (2010) Search and clustering orders of magnitude faster than BLAST. Bioinformatics 26: 2460–2461.

41. Haas BJ, Gevers D, Earl AM, Feldgarden M, Ward DV, et al. (2011) Chimeric 16S rRNA sequence formation and detection in Sanger and 454-pyrosequenced PCR amplicons. Genome Research 21: 494.

42. Caporaso JG, Lauber CL, Walters WA, Berglyons D, Lozupone CA, et al. (2011) Global patterns of 16s rrna diversity at a depth of millions of sequences per sample. Proceedings of the National Academy of Sciences of the United States of America 108 Suppl 1: 4516–4522.

43. Zapala MA, Schork NJ (2006) Multivariate regression analysis of distance matrices for testing associations between gene expression patterns and related variables. Proceedings of the National Academy of Sciences of the United States of America 103: 19430–19435.

44. Chen J, Bittinger K, Charlson ES, Hoffmann C, Lewis J, et al. (2012) Associating microbiome composition with environmental covariates using generalized UniFrac distances. Bioinformatics 28: 2106–2113.

45. Shannon CE (1974) The mathematical theory of communication. 1963: McGraw-Hill. 3–55 p.

46. Segata N, Izard J, Waldron L, Gevers D, Miropolsky L, et al. (2011) Metagenomic biomarker discovery and explanation. Genome Biology 12: 1–18.

47. Langille MG, Zaneveld J, Caporaso JG, Mcdonald D, Knights D, et al. (2013) Predictive functional profiling of microbial communities using 16S rRNA marker gene sequences. Nature Biotechnology 31: 814–821.

48. Caporaso JG, Kuczynski J, Stombaugh J, Bittinger K, Bushman FD, et al. (2010) QIIME allows analysis of high-throughput community sequencing data. Nature methods 7: 335.

49. Martinson VG, Danforth BN, Minckley RL, Rueppell O, Tingek S, et al. (2011) A simple and distinctive microbiota associated with honey bees and bumble bees. Molecular Ecology 20: 619–628.

50. Kwong WK, Moran NA (2016) Gut microbial communities of social bees. Nature Reviews Microbiology.

51. Zheng HQ, Lin ZG, Huang SK, Sohr A, Wu L, et al. (2014) Spore Loads May Not be Used Alone as a Direct Indicator of the Severity of Nosema ceranae Infection in Honey Bees Apis mellifera (Hymenoptera:Apidae). Journal of Economic Entomology 107: 2037–2044.

52. Guo J, Jie W, Chen Y, Evans JD, Dai R, et al. (2015) Characterization of gut bacteria at different developmental stages of Asian honey bees, Apis cerana. Journal of Invertebrate Pathology 127: 110–114.

53. Gilliam M (1997) Identification and roles of non-pathogenic microflora associated with honey bees. FEMS Microbiology Letters 155: 1–10.

54. Su SK, Chen SL, Yu XP, Lin XZ, Hu FL (2001) Isolation and identification of bacteria from pollen and bee bread. Journal of Zhejiang Agricultural University 27: 627–630.

55. Forsgren E, Olofsson TC, Váasquez A, Fries I (2010) Novel lactic acid bacteria inhibiting Paenibacillus larvae in honey bee larvae. Apidologie 41: 99–108.

56. Lourenço AP, Martins JR, Bitondi MMG, Simões ZLP (2009) Trade-off between immune stimulation and expression of storage protein genes. Archives of Insect Biochemistry & Physiology 71: 70–87.

57. Mayack C, Natsopoulou ME, McMahon DP (2015) Nosema ceranae alters a highly conserved hormonal stress pathway in honeybees. Insect Mol Biol 24: 662–670.

58. Elzen PJ, Elzen GW, Lester GE (2004) Compatibility of an organically based insect control program with honey bee (Hymenoptera: Apidae) pollination in cantaloupes. J Econ Entomol 97: 1513–1516.

59. Zheng HQ, Lin ZG, Huang SK, Sohr A, Wu L, et al. (2014) Spore Loads May Not be Used Alone as a Direct Indicator of the Severity of Nosema ceranae Infection in Honey Bees Apis mellifera (Hymenoptera:Apidae). J Econ Entomol 107: 2037–2044.

